# Plant ABCC1 orthologs from agronomically relevant species reveal conserved principles of detoxification transport

**DOI:** 10.64898/2026.09.10.750620

**Authors:** Madushani R.A. Wathukoralage, Alan T. Culbertson

## Abstract

Cytosolic detoxification through vacuolar sequestration is essential for plant growth and stress tolerance. A key driver of this process is the vacuolar ABC transporter, ABCC1, which sequesters diverse amphiphilic anions including glutathione (GS)-conjugates, glucuronic acid-conjugates, and thiol-coordinated metal complexes. However, mechanistic studies of plant ABCC1 have largely relied on Arabidopsis genetics and crude membrane transport assays. Here, phylogenetic and sequence-guided analysis of ABCC1 orthologs from multiple plant species was combined with in vitro functional assays with purified protein to investigate plant ABCC1 transporter activity. Compared to mammalian ABCC1, all plant ABCC1 orthologs contain a conserved bipartite binding pocket with a cationic pocket for anion recognition and a hydrophobic pocket. Key differences between mammalian and plant ABCC1 include a substitution in the cation coordinating histidine residue that is compensated by a peripheral arginine mutation to preserve the cationic electrostatic environment. Furthermore, the plant ABCC1 hydrophobic pocket is enlarged relative to its mammalian counterpart due to several substitutions with smaller side-chain residues. To investigate the in vitro activity of plant ABCC1, several orthologs were screened for expression, and those from maize, date palm, and rice exhibited high biochemical stability and comparable substrate-stimulated ATPase activity. ZmABCC1 was further characterized through a series of assay optimization experiments, which found that its ATPase activity is strongly stimulated by E_2_17βG and GS-conjugates, and inhibited by orthovanadate. This study verifies that ABCC1 is widely conserved across plant species with a consistent substrate binding pocket and activity, supporting its role in cytosolic detoxification in plants.

## Introduction

Plants are sessile organisms that are continually exposed to diverse environmental and anthropogenic toxicants, including heavy metals, xenobiotics, and naturally occurring bioactive compounds. In plants, a common cellular detoxification strategy used is the conjugation of toxic molecules to glutathione (GSH) or glucuronic acid (GlcA), or chelation of heavy metals to phytochelatin (PC), generating derivatives with reduced toxicity (Cobbett and Goldsbrough, 2002). However, even upon conjugation or complex formation, some molecules can retain toxicity in the plant cell cytosol, which they are then further detoxified through sequestration into the plant vacuole, thereby reducing their cytosolic concentration and associated toxicity. This sequestration into the plant vacuole is achieved by the ATP-binding cassette (ABC) transporter ABCC1 (Liu et al., 2001; Lu et al., 1997; Park et al., 2012). More broadly, this ABC transporter-mediated detoxification strategy is conserved across all kingdoms of life, where homologous transporters, including mammalian and yeast ABCC1, either sequester toxic conjugates into intracellular compartments or export them from the cell through broad substrate polyspecificity (Robey et al., 2018), highlighting the conserved role of ABC transporters in cellular detoxification.

ABC transporters are membrane proteins that use energy from ATP to transport molecules across biological membranes. ABC transporters have two transmembrane domains (TMDs) and two nucleotide binding domains (NBDs). ABCC1 (in mammals, plants, and yeast) has an additional TMD, named TMD0, containing five TM helices that is likely involved in protein trafficking or dimerization (Johnson and Chen, 2017; Shinde et al., 2025). Arabidopsis ABCC1 (AtABCC1) was first identified to transport GS-conjugated herbicides (Lu et al., 1997), and has since been shown to have broad substrate specificity. These substrates include GlcA conjugates such as estradiol-17-β-glucuronide (E_2_17βG), GS-conjugates such as S-(2,4-dinitrophenyl)glutathione (DNP-GS), folates, methotrexate (MTX), cyanidin-3-glucoside (C3G), and chlorophyll catabolites (Behrens et al., 2019; Dean et al., 2022; Liu et al., 2001; Lu et al., 1998, 1997; Park et al., 2012). In addition, AtABCC1 has been shown to confer tolerance to heavy metals such as arsenic, mercury, and cadmium, all of which require PCs (Park et al., 2012; Song et al., 2010). AtABCC1 has a functional homolog, AtABCC2, which has overlapping substrate specificity (Liu et al., 2001; Song et al., 2010). This is apparent due to single knock-out plants still having resistance to heavy metals, while double knock-outs have increased sensitivity (Park et al., 2012; Song et al., 2010). However, they do have reported differences in their substrate specificity and transport rates (Liu et al., 2001; Lu et al., 1998). Understanding plant ABCC1 offers numerous potential applications, including engineering herbicide-resistant crops and enhancing plant growth or phytoremediation in heavy metal contaminated soil.

Plant ABCC1 has broad overlapping substrate specificity with mammalian ABCC1, which is a plasma membrane localized multidrug transporter involved in cancer multidrug resistance (Johnson and Chen, 2017). The yeast homolog, Yeast Cadmium Factor 1 (Ycf1), also has similar substrate overlap and is a vacuolar ABC transporter that also transports GS-conjugates or GS-metal complexes into the vacuole (Khandelwal and Tomasiak, 2024). Structural studies of mammalian ABCC1 were first investigated through the bovine ABCC1 structure determined in complex with leukotriene C4 (LTC4), a GS-conjugated epoxide derived from arachidonic acid metabolism, which revealed a bipartite substrate-binding site (Johnson and Chen, 2017). This contains a conserved GSH binding site (P-pocket) along with a hydrophobic pocket (H-pocket). The GSH binding site contains two positions with cationic residues which coordinate the anionic groups on glutathione. Mutational analysis demonstrates that the glutathione coordination residues are critical for function (Shinde et al., 2025; Sun et al., 2025). In addition to GS-conjugates, mammalian ABCC1 exhibits broad polyspecificity for other amphiphilic anions, including GlcA-conjugates (e.g., E_2_17βG) and sulfate-conjugates (e.g., estrone sulfate). Several additional cryo-EM structures demonstrate that substrates bind to a bipartite binding cavity (Sun et al., 2025), establishing the molecular basis for both substrate polyspecificity and synergistic co-transport (Johnson and Chen, 2018; Shinde et al., 2025; Sun et al., 2025; Wang et al., 2020).

In this study, multiple plant ABCC1 homologs were examined to assess conservation within the bipartite substrate-binding site, TMDs, and NBDs of the transporter. Stable homologs suitable for biochemical analysis were identified and each exhibits ATPase activity stimulated by the model ABCC1 substrate, E_2_17βG. Maize ABCC1 (ZmABCC1) was further characterized through assay optimization and determined its substrate specificity. Collectively, these findings provide in vitro functional validation of ABCC1 homologs from agronomically relevant species and offer new insight into the mechanisms of substrate recognition and co-transport in plant ABCC1 transporters.

## Results

### Phylogenetic analysis and conservation of the substrate binding site of plant ABCC1

Arabidopsis ABCC1 has served as a primary model for investigating plant ABCC1 function. To determine the conservation and evolutionary placement of ABCC1 in agronomically relevant species, the complete complement of ABC transporters encoded in eight plant genomes was analyzed, including Arabidopsis, maize, soybean, cowpea, tepary bean, rice, seashore paspalum, and sorghum. Each genome encodes more than 100 ABC transporters, yielding over 1,000 total ABC transporter sequences from these eight species. These were analyzed through phylogenetic analysis together with all human ABC transporters, ABCC1 homologs from yeast and bovine, and additional plant ABCC1 homologs from date palm and saguaro cactus that were identified through NCBI BLAST search.

Phylogenetic analysis of the complete ABC transporter datasets resolved a clearly defined ABCC1 clade (Supplementary Figure 1A) containing at least one ortholog from every species examined (Supplementary Figure 1B), with greater than 70% sequence identity among plant homologs (Figure 1). Arabidopsis contains two closely related paralogs, AtABCC1 and AtABCC2, and soybean was the only other species with more than one ABCC1 homolog (Supplementary Figure 1B). All remaining species encoded a single representative within this clade (Supplementary Figure 1B). Two additional Arabidopsis C-family transporters, AtABCC11 and AtABCC12, formed a neighboring but distinct clade that included related sequences from other species. Although phylogenetically proximal, genetic evidence shows that Arabidopsis abcc1/abcc2 double knock-out mutants are hypersensitive to heavy metals despite the presence of ABCC11 and ABCC12 (Park et al., 2012; Song et al., 2010), indicating functional distinction from ABCC1. Accordingly, all subsequent analyses were restricted to transporters within the ABCC1 orthologous clade (Figure 1 and Supplementary Figure 1B).

**Figure 1.**
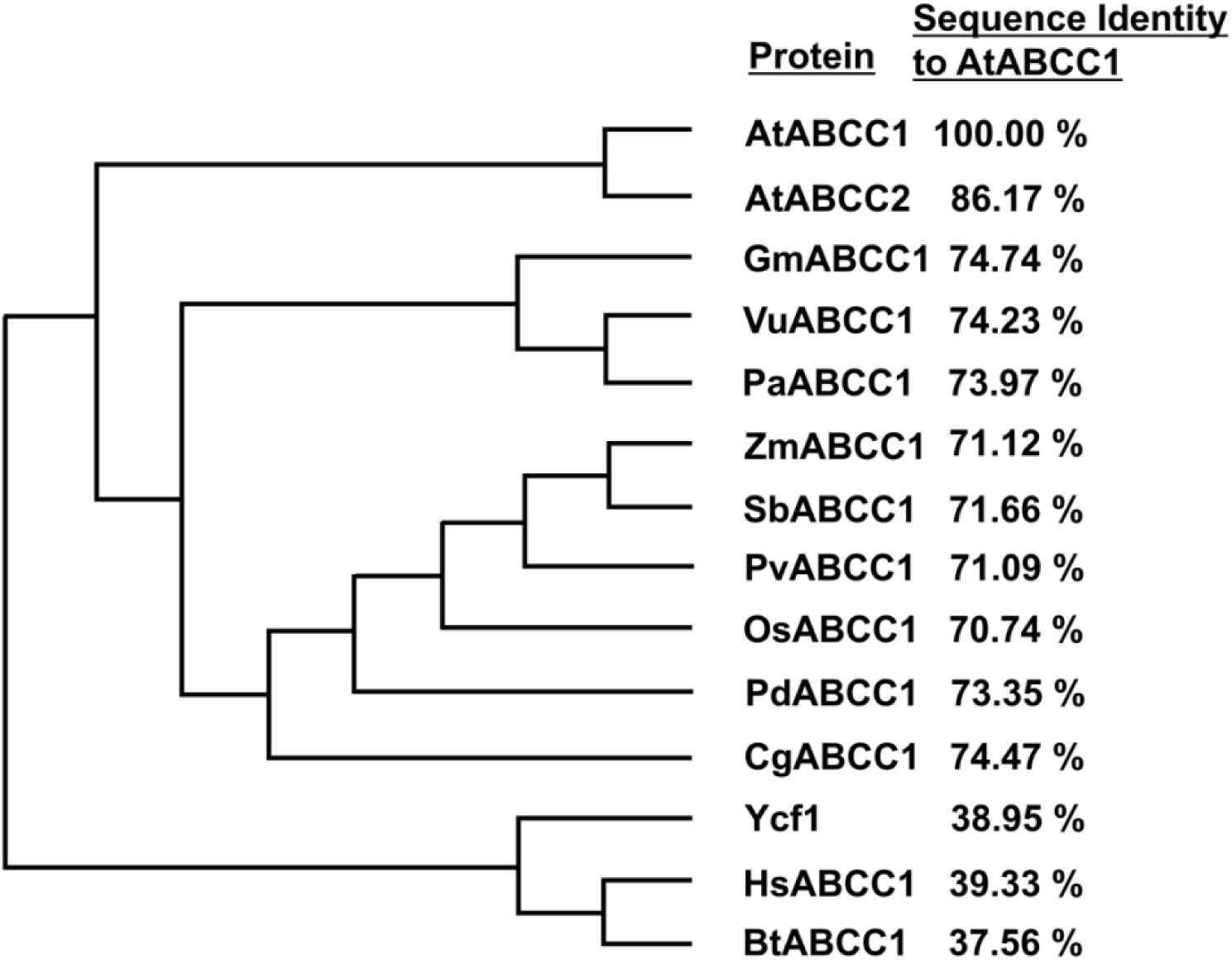
Phylogenetic analysis of ABCC1 homologs. Phylogenetic tree of ABCC1 homologs. Plants included are Arabidopsis (At, *Arabidopsis thaliana*), Soybean (Gm, *Glycine max*), Cowpea (Vu, *Vigna unguiculata*), Tepary bean (Pa, *Phaseolus acutifolius*), Maize (Zm, *Zea* mays), Sorghum (Sb, *Sorghum bicolor*), Seashore paspalum (Pv, *Paspalum vaginatum*), Rice (Os, *Oryza sativa*), Date palm (Pd, *Phoenix dactylifera*), and Saguaro cactus (Cg, *Carnegiea gigantea*). Also included are mammalian homologs of ABCC1 from Human (Hs, *Homo sapiens*) and Bovine (Bt, *Bovine taurus*), and the Yeast homolog Ycf1 (*Saccharomyces cerevisiae*). Next to phylogenetic tree are percent sequence identity relative to AtABCC1.

In addition to comparing full-length sequence identity (Figure 1), pairwise percent sequence identity was quantified for individual domains, including TMD0, TMD1, TMD2, NBD1, NBD2, the regulatory R-domain (located between NBD1 and TMD2), and the C-terminal domain (Figure 2A). Across plant ABCC1 homologs, TMD1, TMD2, NBD1, and NBD2 exhibited consistently high sequence conservation, with percent identities all being greater than 75% relative to AtABCC1. In contrast, the C-terminal domain displayed moderate conservation (≥63% identity across plant homologs), whereas TMD0 showed lower and more variable identity (approximately 50–75%; Figure 2A). The R-domain also exhibited comparatively low sequence conservation. Overall, the strong conservation of TMD1-2 and NBD1-2 supports the conclusion that these core structural and catalytic domains are under substantial evolutionary constraint. In contrast, the greater variability observed in TMD0, the R-domain, and the C-terminal region suggests comparatively relaxed sequence constraint, potentially reflecting regulatory or lineage-specific divergence rather than essential functional roles. As expected, given their lower overall sequence identity relative to plants, the mammalian and yeast orthologs exhibited reduced sequence conservation across all domains (Figure 2A). Further, sequence conservation of functionally important motifs within the NBDs was analyzed, including Walker A, Walker B, Signature, A-loop, Q-loop, and H-switch (Figure 2B-C). In mammalian and yeast ABCC1, nucleotide binding site (NBS1) is degenerate due to substitutions in ATP binding motifs, including Walker B and Signature Motif, that make it catalytically inactive (Johnson and Chen, 2018). This character is conserved in plant ABCC1, all of which contain a degenerate NBS1 and canonical NBS2 (Figure 2C).

**Figure 2.**
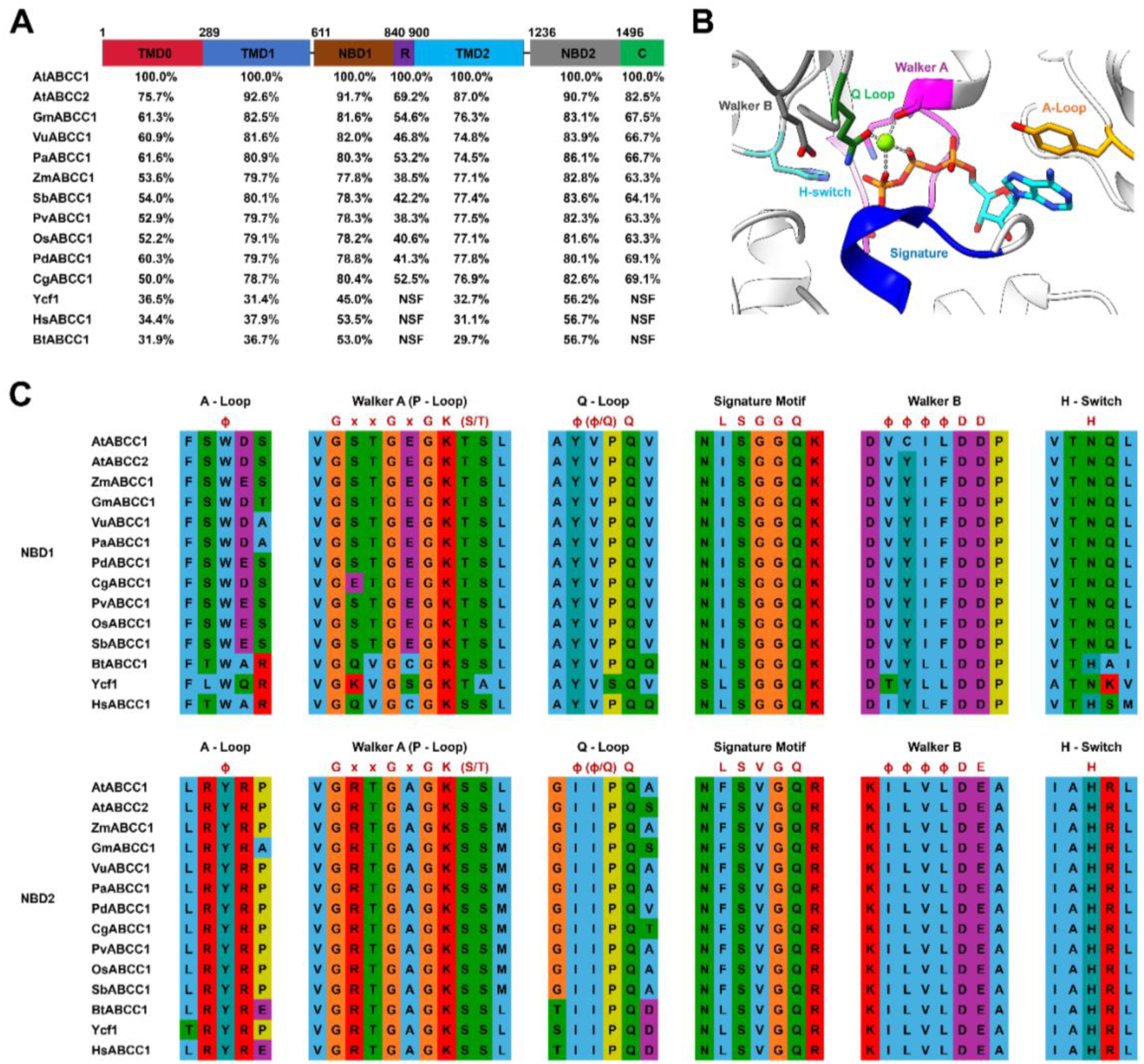
Sequence conservation of the domains of ABCC1 and the substrate binding site and nucleotide binding site architecture of ABCC1. **A.** Percent sequence identities of each domain in ABCC1 homologs compared to AtABCC1 (TMD: transmembrane domain; NBD: nucleotide binding domain; R: regulatory domain; C: C-terminus). NSF indicates NCBI pairwise sequence alignments resulted in “no significant similarity found”. **B.** Mg^2+^/ATP bound nucleotide binding site (NBS) of BtABCC1. A-Loop, Walker A, Q-Loop, Signature motif, Walker B and H-Switch of nucleotide binding site are indicated in respective colors as in the labels. **C.** Sequence conservation of six core nucleotide-binding motifs in nucleotide binding-domains (NBDs) of ABCC1 homologs. A Nucleotide binding site (NBS) is formed by the A-loop, Walker A, Walker B, Q-loop, and H-switch on one NBD, together with the signature motif on the other NBD. Accordingly, NBS2 has all the conserved motifs, whereas NBS1 contains several substitutions, including alterations in the signature motif and Walker B, rendering NBS1 a degenerate site.

Sequence conservation within the substrate-binding site of plant ABCC1 homologs was compared using the cryo-EM structure of bovine ABCC1 (BtABCC1) bound to LTC4 as a reference (Johnson and Chen, 2017). Although mammalian ABCC1 shares approximately 35% sequence identity with plant ABCC1 (Figure 1), there is substantial overlap in substrate specificity, including glutathione conjugates, glucuronic acid conjugates, and heavy metal complexes (Liu et al., 2001; Song et al., 2010; Sun et al., 2025). The BtABCC1-LTC4 structure provides a useful model because LTC4 contains a GSH moiety conjugated to leukotriene A4, enabling comparison with GS-conjugated substrates transported by plant ABCC1. The structure reveals a bipartite binding pocket consisting of a positively charged region (P-pocket) and a hydrophobic region (H-pocket) (Johnson and Chen, 2017). The P-pocket (shown as red circles in Figure 3) contains two primary binding sites for the carboxylate groups of the GSH moiety (Figure 3). In BtABCC1, one site is formed by Arg1196 and Arg1248, and the second by Lys332 and His335. The arginine and lysine residues are conserved in plant ABCC1 homologs (Lys309, Arg1136, and Arg1193 in AtABCC1), whereas His335 is replaced by asparagine in all plant sequences (Asn312 in AtABCC1), removing a potential electrostatic interaction (Figure 3). This substitution may be functionally compensated by a concurrent Gln450-to-Arg replacement observed in plants (Arg426 in AtABCC1). Although Gln450 in BtABCC1 does not directly contact the LTC4 carboxylate (approximately 6 Å distance), substitution to arginine in plant homologs would likely allow extension to permit formation of a stabilizing electrostatic interaction. Thus, this analysis supports that these anion binding sites are conserved in plant ABCC1 orthologs (Figure 3).

**Figure 3.**
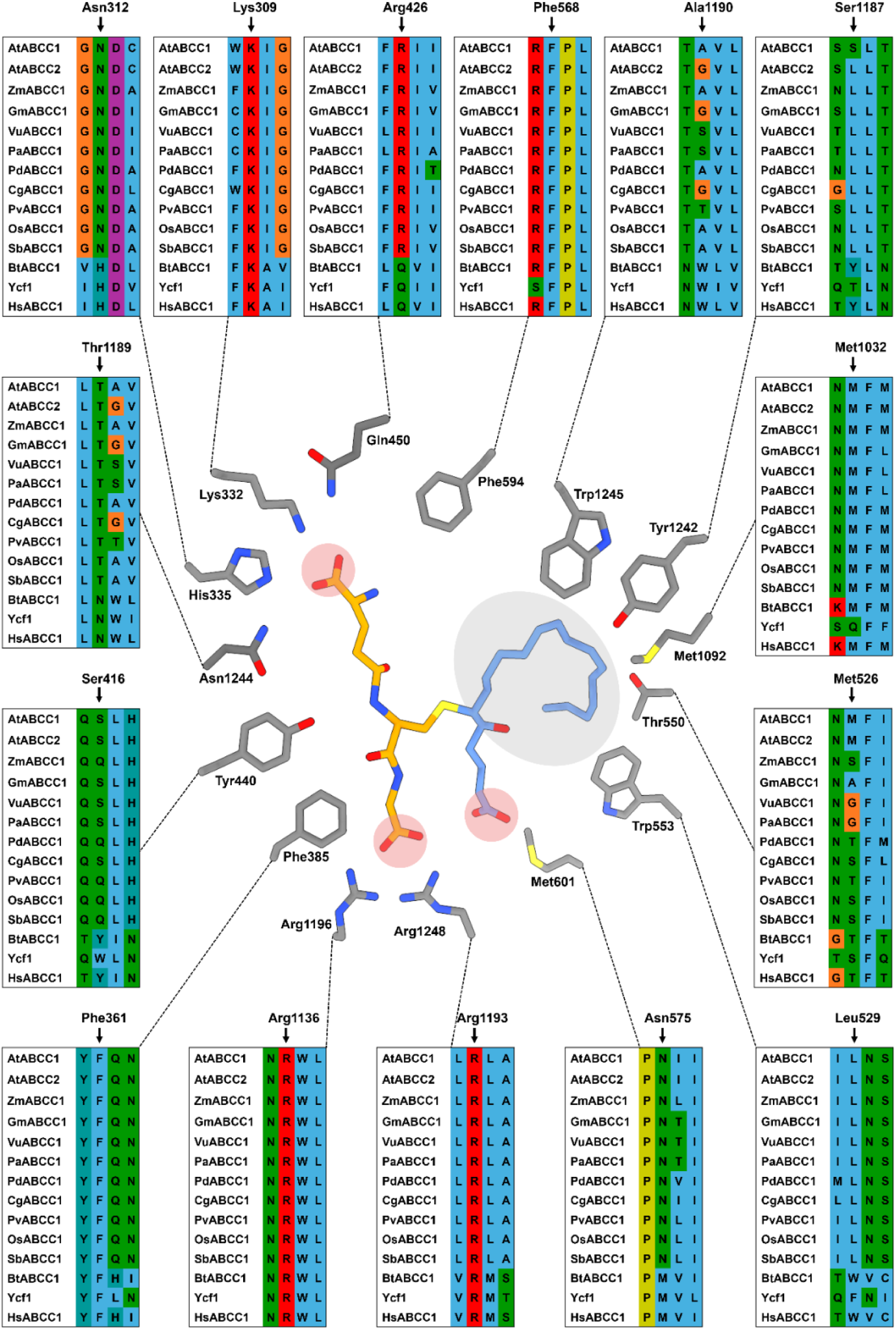
Sequence conservation of the substrate-binding site of plant ABCC1. Sequence alignments of residues involved in substrate binding of ABCC1, defined based on the cryo-EM structure of the leukotriene C4 (LTC4) bound to bovine ABCC1 (PDB 5UJA). Residues shown in the central panel of the figure correspond to those within the binding site of bovine ABCC1. Sequence alignments shown surrounding the structure compare each bovine binding-site residue to the corresponding positions in ABCC1 homologs. Residue numbers adjacent to amino acids in the central panel correspond to bovine ABCC1 numbering, and residue numbers above each sequence alignment indicate the corresponding position in Arabidopsis ABCC1.

In contrast to the P-pocket, the H-pocket exhibits lower sequence conservation relative to mammalian ABCC1. Among the residues analyzed, only Met1092 and Phe594 are conserved across all plant species examined (Figure 3). Although several H-pocket residues differ between plant and animal orthologs, many substitutions retain hydrophobic character, thus retaining the bipartite nature of the ABCC1 binding pocket. Notably, three bulky hydrophobic residues in mammalian ABCC1 are replaced by smaller hydrophobic residues in plant homologs, including Trp553 to Leu (Leu529 in AtABCC1), Tyr1242 to Ser/Leu (Ser1187 in AtABCC1), and Trp1245 to Ala (Ala1190 in AtABCC1) (Figure 3). These substitutions reduce side-chain volume and are likely to increase the accessible surface area of the H-pocket, resulting in a larger hydrophobic cavity in plant ABCC1 homologs compared to their mammalian counterparts. In summary, although human and bovine ABCC1 share only approximately 35% sequence identity with AtABCC1, the majority of key residues within the P-pocket are conserved, and the hydrophobic character of the H-pocket is largely maintained. These observations indicate that the bipartite organization of the substrate-binding site is preserved in plant ABCC1, consistent with that observed in mammalian orthologs, and support the conservation of core functional features across species.

### Protein expression screening of plant ABCC1 orthologs

To experimentally verify the overlapping functions of these plant ABCC1 homologs, stable orthologs suitable for *in vitro* characterization were screened. This was achieved by transient protein expression of all ABCC1 orthologs in HEK293F cells followed by fluorescence-detection size-exclusion chromatography (FSEC) to assess protein expression levels. All genes were synthesized and cloned into pFastBacMam with a C-terminal GFP tag for expression screening. ZmABCC1, PdABCC1, and OsABCC1 exhibited the highest expression levels with minimal degradation or aggregation (Figure 4A and Supplementary Figure 2A). These three homologs were subsequently subcloned into pFastBac1 with a C-terminal maltose-binding protein (MBP) tag and expressed in insect cells using the baculovirus expression system. All proteins were purified with the MBP tag to greater than 90% purity (Supplementary Figure 2B). Functional activity was assessed using an ATP hydrolysis (ATPase) assay. Substrate-stimulated ATP hydrolysis assays offer a validated, high-throughput approach to screen compounds for transporter activity. While this assay measures ATP hydrolysis rather than direct substrate transport, ATPase activity typically correlates directly with substrate transport. For example, studies on mammalian ABCC1 demonstrate a strong correlation between ATP hydrolysis and substrate transport, yielding consistent kinetic parameters across both assay types (Loe et al., 1996; Mao et al., 1999). Assays were conducted in triplicate with 50 µL reactions within a 96-well plate using purified ZmABCC1 (Figure 4D), incubated at 37 °C for one hour, and terminated with SDS before adding molybdate for color development.

**Figure 4.**
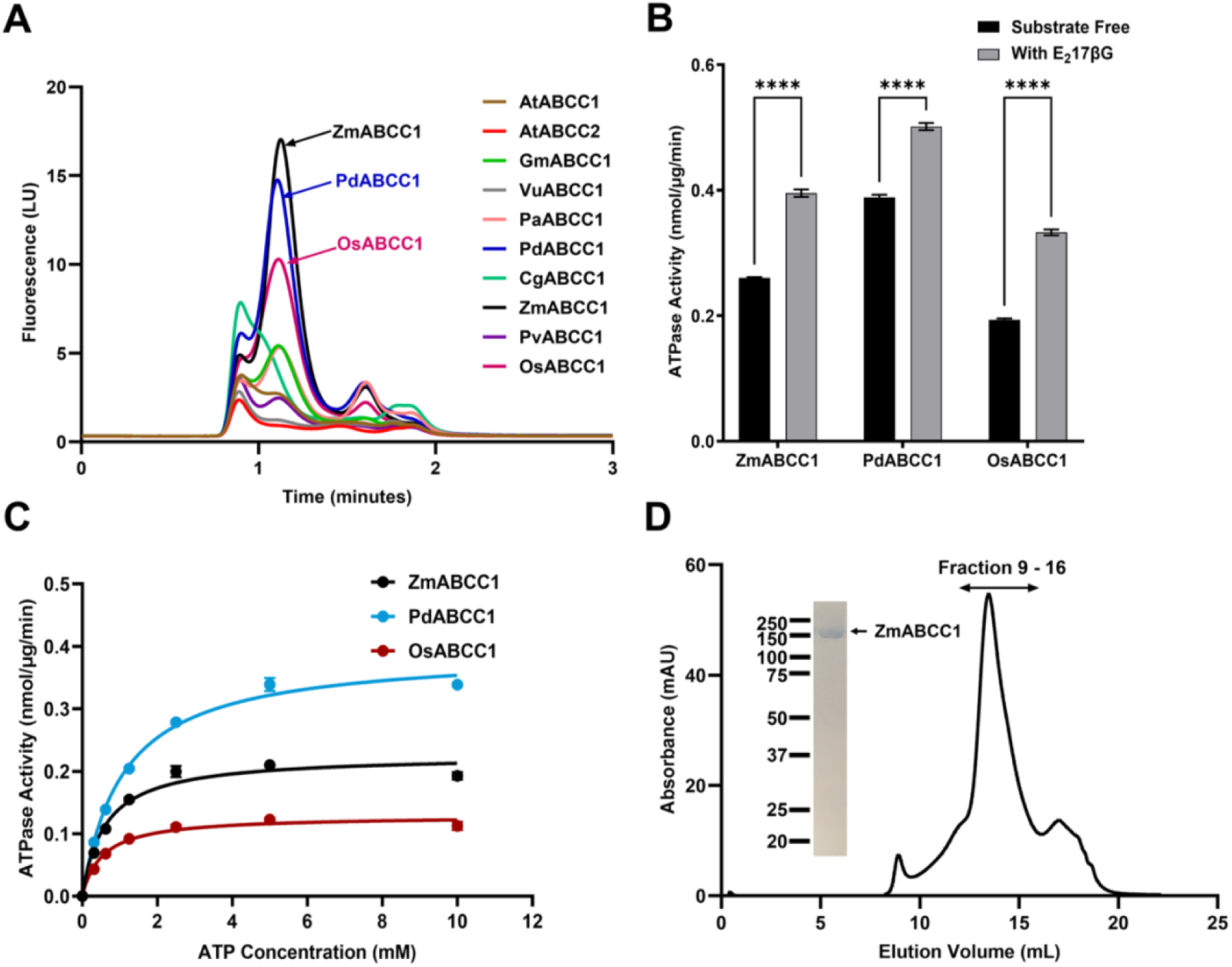
Functional characterization of high expressing ABCC1 orthologs. **A.** Fluorescent size exclusion chromatography (FSEC) chromatograms of GFP-tagged ABCC1 homologs from nine plant species. **B.** ATPase activity of purified C-terminal MBP-tagged ZmABCC1, PdABCC1 and OsABCC1. Each protein has a basal ATPase activity (Substrate free), and the activity was stimulated in the presence of model substrate E_2_17βG (With E_2_17βG). **C.** ATP saturation analyses of purified C-terminal MBP-tagged ZmABCC1 (K_m_ = 0.61 mM, V_max_ = 0.23 nmol µg^-1^ min^-1^), PdABCC1 (K_m_ = 1.09 mM, V_max_ = 0.39 nmol µg^-1^ min^-1^), and OsABCC1 (K_m_ = 0.54 mM, V_max_ = 0.13 nmol µg^-1^ min^-1^). **D.** Size exclusion chromatogram of ZmABCC1 following MBP cleavage. The corresponding SDS-PAGE gel of pooled fractions confirms the presence of highly purified ZmABCC1. Data in B and C are presented as mean *±* standard deviation from triplicates (n=3). Statistical significance of the data in B was determined by two-way ANOVA followed by Sidak multiple comparison test (ns, P ≥ 0.05 (not statistically significant); *, P < 0.05; **, P < 0.01; ***, P < 0.001; ****, *P* < 0.0001).

Each plant ortholog displayed basal ATPase activity in the presence of ATP, and ATPase activity was further stimulated by the model ABCC1 substrate E_2_17βG, a validated substrate of AtABCC1 (Figure 4B). Furthermore, because plant ABCC1 has been shown to transport E_2_17βG (Liu et al., 2001), these data support that substrate-stimulated ATPase is a reliable method for screening transport activity. ATP saturation analysis yielded apparent K_m_ values for ATP ranging from 0.5 to 1.0 mM (Figure 4C), consistent with reported values for Arabidopsis ABCC2 (K_m_ ≈ 0.4 mM) (Qiu et al., 2025) and mammalian ABCC1 (K_m_ ≈ 0.5 mM) (Johnson and Chen, 2018). Overall, the high sequence identity (Figure 1), conservation of the substrate-binding site (Figure 3), and comparable in vitro functional activity (Figure 4B-C) support that ZmABCC1, PdABCC1, and OsABCC1 are functional orthologs of AtABCC1 with conserved biochemical properties.

### Characterization of substrate specificity of ZmABCC1

Due to its high biochemical stability, ZmABCC1 was selected for detailed characterization. ZmABCC1 was expressed in Sf9 cells using the baculovirus expression system and purified via a C-terminal maltose-binding protein (MBP) affinity tag. The MBP tag was removed by TEV protease cleavage, and the protein was further purified by size-exclusion chromatography (SEC), yielding a single, symmetrical peak corresponding to monodisperse ZmABCC1 (Figure 4D). Peak fractions were analyzed by SDS-PAGE gel, which showed a predominant band at approximately 165 kDa, consistent with highly purified, full-length ZmABCC1 lacking the affinity tag (Figure 4D).

The functional activity of purified ZmABCC1 was evaluated using ATPase assays. As expected, ZmABCC1 displayed basal ATPase activity which was stimulated by roughly 45% with the addition of the model substrate E_2_17βG (Figure 5A). This activity was abolished with the addition of sodium orthovanadate, confirming inhibition of ABC transporter-mediated ATP hydrolysis (Figure 5B). Assay conditions were optimized, demonstrating that activity increased with protein concentration (Figure 5C), remained linear for up to 2 hours (Figure 5D), and was maximal at 37 °C (Figure 5E). ATP saturation analysis of MBP-cleaved ZmABCC1 (Figure 5F) yielded kinetic parameters comparable to those obtained prior to cleavage (Figure 4C), indicating that the affinity tag did not affect protein function.

**Figure 5.**
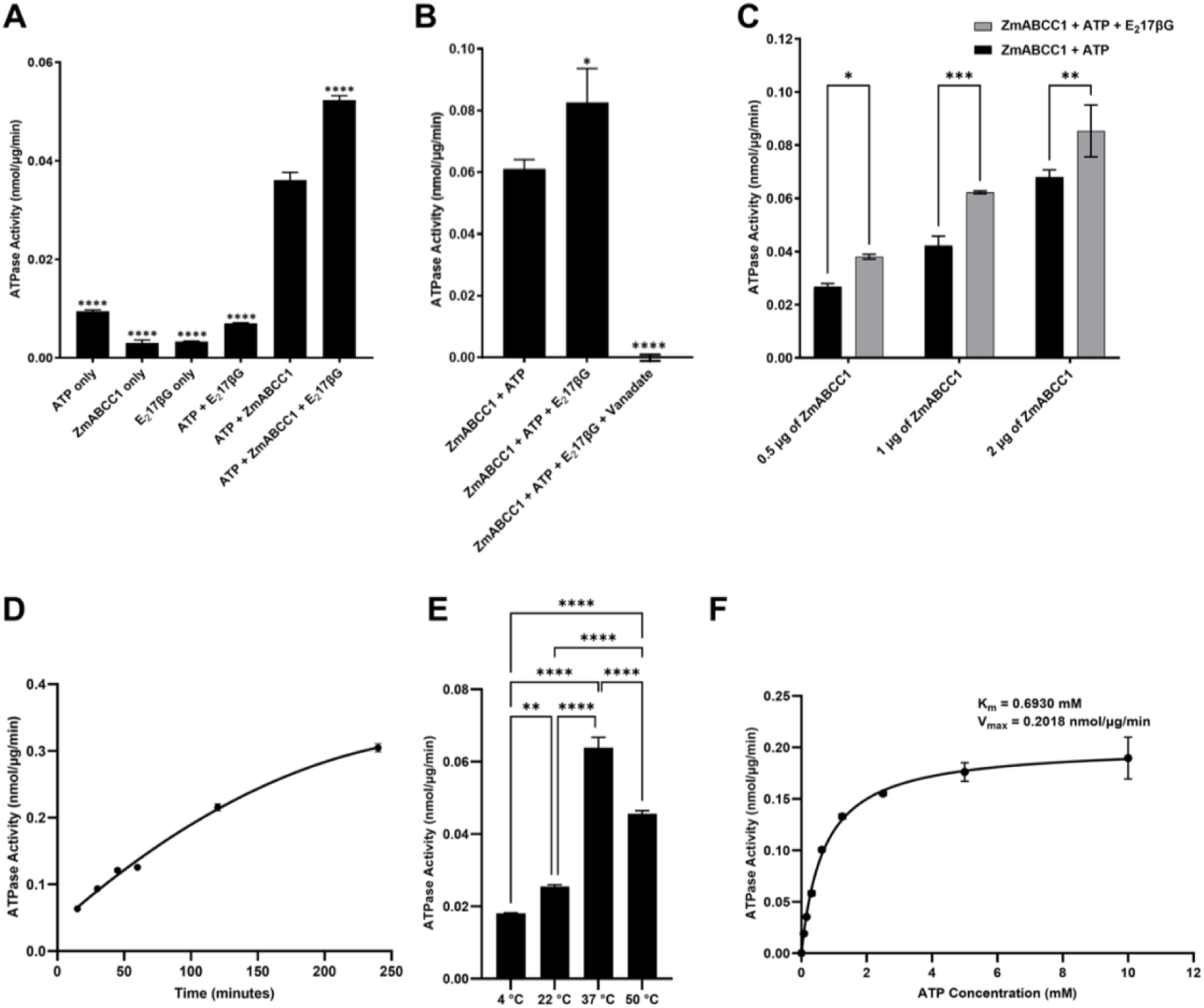
Characterization and optimization of ZmABCC1 ATPase activity and substrate screening of ZmABCC1. **A.** ATPase activity of purified ZmABCC1 conducted with the control experiments for the verification of the in vitro ATPase activity. All reactions contained identical buffers but with additional of only selected components including ATP alone, protein alone, substrate alone, and combinations of these. **B.** Inhibition of ZmABCC1 ATPase activity in the presence of 2 mM sodium orthovanadate. **C-E.** Optimization of the assay conditions, including protein concentration (**C**), incubation time (**D**), and reaction temperature (**E**). **F.** ATP saturation kinetics of ZmABCC1 with cleaved MBP (K_m_ = 0.693 mM, V_max_ = 0.202 nmol µg^-1^ min^-1^). All data is presented as mean *±* standard deviation from triplicates (n=3). Statistical significance of the data in A and B was determined by one-way ANOVA followed by Dunnett’s multiple comparison test, in C was determined by two-way ANOVA followed by Sidak multiple comparison test, and in E was determined by one-way ANOVA followed by Tukey’s multiple comparison test (ns, P ≥ 0.05 (not statistically significant); *, P < 0.05; **, P < 0.01; ***, P < 0.001; ****, *P* < 0.0001).

### Substrate screen of substrate-stimulated ATPase activity

AtABCC1 exhibits broad substrate polyspecificity, transporting a diverse range of anionic compounds. To assess whether this property is conserved in maize ABCC1, purified ZmABCC1 was evaluated using a panel of proposed plant and mammalian ABCC1 substrates, including anionic small molecules, glutathione conjugates, and glucuronic acid conjugates. First, verified plant ABCC1 substrates were selected, including E_2_17βG, GSH, folate, methotrexate (MTX), and cyanidin-3-glucoside (C3G) (Behrens et al., 2019; Dean et al., 2022; Liu et al., 2001; Raichaudhuri et al., 2009). The known mammalian ABCC1 substrate including 4-hydroxy-nonenal-glutathione (HNE-GS), estrone-3-sulfate (E3S), calcein, and fluorescein were also included (Deeley and Cole, 2006). In addition, γ-glutamyl-cysteine (γ-Glu-Cys) as a structural analog of GSH and 4-methylumbelliferyl-β-D-glucuronide (4-MUG) as a model glucuronide conjugate were included to define the minimal chemical determinants required for recognition and transport. Among the compounds tested, HNE-GS, E3S, and E_2_17βG produced robust stimulation of ATPase activity (Figure 6A). In contrast, folate, GSH, γ-Glu-Cys, MTX, 4-MUG, calcein, and fluorescein resulted in minimal or no stimulation (Figure 6A). Notably, C3G, previously proposed as an AtABCC1 substrate (Dean et al., 2022), reduced ATPase activity under the conditions tested (Figure 6A). These results were reproducible with biological replicates from completely different protein purification preparations (Supplementary Figure 3B-C).

**Figure 6.**
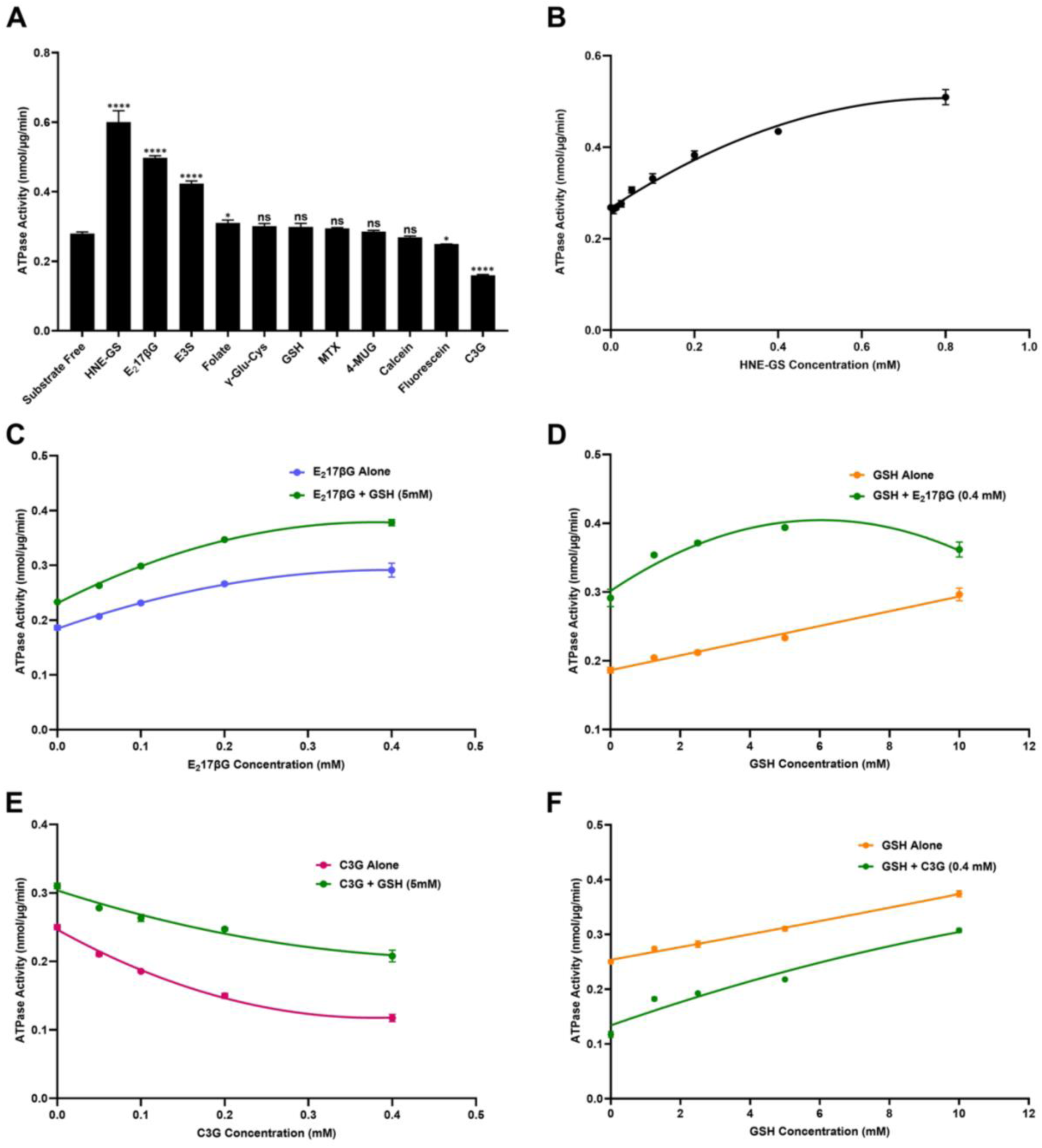
Substrate screening and synergistic ATPase activity of ZmABCC1 with glutathione. **A.** Substrate screening of purified ZmABCC1. All the substrates were tested at final concentration of 0.4 mM unless indicated otherwise. Methotrexate (MTX) was tested at 0.2 mM. Substrates included 4-hydroxy-nonenal-glutathione (HNE-GS), estradiol 17β-D-glucuronide (E_2_17βG), estrone-3-sulfate (E3S), Folate, γ-L-glutamyl-L-cysteine (γ-Glu-Cys), reduced L-glutathione (GSH), MTX, 4-methylumbelliferyl-β-D-glucuronide (4-MUG), Calcein, Fluorescein, and cyanidin-3-glucoside (C3G). **B**. ATPase activity of ZmABCC1 with increasing concentrations of HNE-GS up to 0.8 mM. **C.** ATPase activity of purified ZmABCC1 with increasing concentrations of E_2_17βG up to 0.4 mM in the presence or absence of 5 mM GSH. **D.** ATPase activity of purified ZmABCC1 with increasing concentrations of GSH up to 10 mM in the presence or absence of 0.4 mM E_2_17βG. **E.** ATPase activity of purified ZmABCC1 with increasing concentrations of C3G up to 0.4 mM in the presence or absence of 5 mM GSH. **F.** ATPase activity of purified ZmABCC1 with increasing concentrations of GSH up to 10 mM in the presence or absence of 0.4 mM C3G. All data is presented as mean ± standard deviation from triplicates (n=3). Statistical significance of the data in A was determined by one-way ANOVA followed by Dunnett’s multiple comparison test (ns, P *≥* 0.05 (not statistically significant); *, P < 0.05; **, P < 0.01; ***, P < 0.001; ****, P < 0.0001).

### Kinetic analysis of ZmABCC1 substrates and co-transport with GSH

To further assess the specificity of different substrates for ZmABCC1 substrate-stimulated ATPase activity, saturation curves of a selected set of substrates were performed. The substrates chosen for these assays were a representative GSH- and GlcA-conjugate (HNE-GS and E_2_17βG), GSH alone, C3G, and folate. In addition, because both plant and mammalian ABCC1 transporters have been reported to exhibit synergistic substrate transport in the presence of GSH (Liu et al., 2001; Rothnie et al., 2008), substrate saturation experiments of E_2_17βG and C3G in the presence of 5 mM GSH were performed. E_2_17βG was selected because it produced the strongest substrate-stimulated ATPase activity among compounds not pre-conjugated to GSH (Figure 6A). C3G was selected because it has been reported as an AtABCC1 substrate (Behrens et al., 2019), yet under assay conditions in this study, C3G inhibited ATPase activity in the absence of GSH (Figure 6A).

HNE-GS showed increased substrate stimulated ATPase with increasing substrate concentration, producing a K_m_ of 0.55 mM, but did not reach full saturating activity within the conditions tested (Figure 6B and Supplementary Figure 3D). E_2_17βG saturation curve produced a K_m_ of 0.32 mM (Supplementary Figure 3E). Saturation curve of E_2_17βG in the presence of 5 mM GSH followed similar saturation kinetics to E_2_17βG alone, while the inclusion of GSH increased the maximal ATPase activity, and the magnitude of stimulation increased with higher E_2_17βG concentrations (Figure 6C), consistent with a synergistic interaction (Liu et al., 2001). Reciprocal saturation experiments were performed by varying GSH concentration in the presence or absence of E_2_17βG (Figure 6D). In the absence of E_2_17βG, GSH-dependent ATPase activity increased linearly, suggesting weak or low-affinity interaction when alone. In contrast, in the presence of E_2_17βG, GSH produced a saturable response, indicating that E_2_17βG enhances the apparent affinity of ZmABCC1 for GSH (Figure 6D). These results support cooperative interactions between E_2_17βG and GSH, as reported before (Liu et al., 2001).

Analogous co-transport experiments were performed with C3G. Increasing C3G concentrations produced dose-dependent inhibition of ATPase activity (K_i_ = 0.21 mM; Supplementary Figure 3F), reaching a maximal effect at higher concentrations, consistent with saturable inhibition (Figure 6E). In the presence of 5 mM GSH, increasing C3G still inhibited ATPase, but the inhibitory effect of C3G was marginally attenuated, as indicated by a slightly reduced decline in ATPase activity with increasing C3G concentration (Figure 6E). When GSH concentration was varied in the absence or presence of C3G (Figure 6F), ATPase activity remained lower in the presence of C3G across all GSH concentrations, although activity increased with increasing GSH. Unlike the E_2_17βG condition, GSH did not exhibit clear saturation behavior in the presence of C3G. Together, these findings indicate that E_2_17βG and C3G differentially influence the interaction of ZmABCC1 with GSH, consistent with distinct substrate-binding and transport mechanisms. Further, because folate is proposed to be a substrate for AtABCC1 (Raichaudhuri et al., 2009), a similar saturation curve was performed, but observed no change in ATPase with increasing folate concentration (Supplementary Figure 3A).

## Discussion

Plant ABCC1 has numerous physiological functions including chlorophyll catabolism, xenobiotic detoxification, and the storage of cell pigmentation and antimicrobial compounds (Behrens et al., 2019; Dean et al., 2022; Lu et al., 1998; Song et al., 2010). Due to these diverse functions in plants, ABCC1 has potential for multiple applications in agriculture and biotechnology because it mediates vacuolar sequestration of chemically diverse toxins. One potential application is the engineering of herbicide resistance by designing herbicides that are efficiently transported by ABCC1 and deploying these compounds in crop lines that overexpress ABCC1. This is supported by evidence that AtABCC1 and AtABCC2 can transport glutathione-conjugated herbicides, such as metolachlor-GS (Frelet-Barrand et al., 2008; Lu et al., 1998, 1997). However, single knockout mutants in AtABCC2 exhibited only minor effects on herbicide sensitivity, with little to no phenotypic changes observed for AtABCC1 knock-outs (Frelet-Barrand et al., 2008), but abcc1/abcc2 double mutants were not evaluated for herbicide resistance. Studies of heavy metal detoxification have shown that plant sensitivity to heavy metals becomes most evident primarily when both genes are disrupted, consistent with their functional redundancy (Song et al., 2010). Therefore, it remains possible that the limited phenotypes observed in single mutants reflect overlapping activity between these transporters. However, even if applicable, binding site conservation analysis indicates that the substrate-binding pocket is highly conserved across plant ABCC1 orthologs (Figure 3), suggesting similar substrate preferences among diverse species. As a result, simple overexpression of a native ABCC1 may provide limited durability, because weeds that possess conserved ABCC1 homologs could be selected for enhanced expression, thereby acquiring comparable detoxification capacity under herbicide pressure. Achieving long-term resistance would therefore likely require engineering the ABCC1 substrate-binding pocket to alter substrate specificity toward compounds that are not efficiently recognized by endogenous weed ABCC1 transporters.

Despite these potential applications, most mechanistic and functional studies have focused on Arabidopsis ABCC1, limiting translation to agronomically relevant species. Here, this was expanded on by performing phylogenetic and sequence analyses of plant ABCC1 orthologs and by biochemical characterization of agronomically relevant orthologs. The sequence analyses indicate that ABCC1 is broadly conserved across the plant species examined, with each plant analyzed having at least one ortholog with sequence identity greater than 70% (Figure 1), consistent with an essential role in detoxification. Phylogenetic analysis indicates that most species encode a single ABCC1 ortholog, whereas Arabidopsis and soybean encode two, suggesting lineage-specific duplication events and potential functional redundancy or specialization in these species (Supplementary Figure 1). The core architecture, including the TMDs and NBDs, is highly conserved among orthologs (Figure 2A) and all contain a degenerate nucleotide binding site (Figure 2C), similar to mammalian ABCC1 (Khandelwal and Tomasiak, 2024).

At the substrate-binding site, plant ABCC1 homologs retain the bipartite organization described for mammalian ABCC1, including cationic features that coordinate acidic groups on GSH (Figure 3). Notably, one anion binding site in mammalian ABCC1 includes a histidine-lysine pair, whereas plant homologs lack the histidine and instead exhibit a conserved substitution at another proximal residue (Gln-to-Arg) consistent with maintenance of electrostatic recognition by an alternative basic residue (Figure 3). Mutational analyses of mammalian and yeast ABCC1, as well as functional characterizations of plant homologs, highlight the conserved importance of the basic P-pocket in substrate recognition (Haimeur et al., 2002; Shinde et al., 2025; Situ et al., 2004; Sun et al., 2025). Positively charged residues coordinating the anionic groups of glutathione, including Lys309, Arg1136, and Arg1193 in Arabidopsis ABCC1 (corresponding to Lys332, Arg1196, and Arg1248 in bovine ABCC1), are required for transport function (Haimeur et al., 2002; Qiu et al., 2025; Situ et al., 2004). Mutations at these positions in both mammalian and plant orthologs severely reduce substrate-stimulated ATPase activity and direct substrate transport, supporting a conserved evolutionary role for electrostatic recognition within the P-pocket.

The H-pocket also remains predominantly hydrophobic in plant homologs, but several positions contain smaller side chains relative to mammalian ABCC1, consistent with a larger hydrophobic cavity. Bulky aromatic side chains present in mammalian ABCC1, including Tyr440, Tyr1242, and Trp1245, are replaced in plant orthologs by smaller residues, specifically Ser416, Ser1187 (or Leu), and Ala1190, respectively (Figure 3). In mammalian ABCC1, mutational substitution of these aromatic positions with smaller or non-aromatic residues impairs substrate transport (Grant et al., 2008; Ito et al., 2001; Sun et al., 2025), suggesting fundamental differences in the mode of substrate binding between plant vs mammalian or yeast ABCC1. These structural differences, supported by prior mammalian mutagenesis studies, indicate that plant ABCC1 possesses an expanded H-pocket, potentially capable of accommodating a broader range of hydrophobic moieties. Further comparative studies are needed to confirm these structural observations and define their functional consequences.

Substrate screening of purified ZmABCC1 aligns with established models of ABCC1 function, demonstrating robust stimulation of ATPase activity by the glutathione conjugate HNE-GS and the glucuronide conjugate E_2_17βG. Furthermore, E_2_17βG exhibits a cooperative, synergistic activation with GSH, consistent with co-transport mechanisms reported for mammalian and plant orthologs. Unexpectedly, folate, MTX, and C3G failed to stimulate ZmABCC1 ATPase activity, contrasting with previous studies with AtABCC1 (Dean et al., 2022; Raichaudhuri et al., 2009). Notably, C3G produced a dose-dependent reduction in ATPase activity, suggesting a potential inhibitory interaction. Because prior studies demonstrating C3G transport by AtABCC1 included 5 mM GSH in their assays, C3G kinetics in the presence of constant 5 mM GSH were evaluated. Under these conditions, C3G maintained an inhibitory profile, although the relative magnitude of ATPase suppression was attenuated. Furthermore, reciprocal saturation curve demonstrated that the presence of C3G did not produce a saturable response of ZmABCC1 ATPase with GSH, suggesting that C3G and E_2_17βG engage the substrate-binding cavity through distinct mechanisms. Structural investigations will be required to define the exact molecular mechanisms underlying GSH synergy between E_2_17βG and C3G.

AtABCC2 is reported to exhibit higher transport activity than AtABCC1 for multiple substrates, including E_2_17βG, DNP-GS, and PC-metal complexes (Liu et al., 2001; Lu et al., 1998; Song et al., 2010). Despite reported functional differences between AtABCC1 and AtABCC2, sequence comparison of the substrate-binding pocket reveals only limited variation, which changes only at Ala1190 and Ser1187 of AtABCC1, whereas AtABCC2 contains Gly and Leu at the corresponding positions (Figure 3). Position 1190 is variable among orthologs, while position 1187 in most ABCC1 orthologs more closely resembles AtABCC2 than AtABCC1 with a Leucine (Figure 3). Thus, differences in activity between AtABCC1 and AtABCC2 may be influenced by these substitutions within the substrate-binding pocket. However, these changes are relatively minor within its binding site, especially within a transporter that accommodates such a broad range of chemical structures such as ABCC1. Therefore, the observed functional differences may also arise from other factors, such as variations along the substrate translocation pathway or differences in coupling between substrate binding and conformational transitions during the transport cycle.

To date, functional characterization of non-Arabidopsis ABCC1 orthologs remains limited to two studies, including a knockout analysis of rice ABCC1 and expression profiling in maize under cadmium stress (Qu et al., 2023; Song et al., 2014). Under heavy metal stress, ZmABCC1 expression is significantly upregulated in roots of Cd-tolerant maize cultivars (Qu et al., 2023). In vitro characterization of ZmABCC1 in this study confirms that its substrate-stimulated ATPase activity and kinetic behavior closely mirror those of orthologs such as AtABCC1, establishing a conserved mechanism of vacuolar transport across plant species. Furthermore, screening of ZmABCC1 substrates determines the broad substrate specificity of plant ABCC1. Characterizing these specificities in the maize ABCC1, along with documenting the conservation of the protein and binding site amino acids, provides stronger translational potential for agricultural applications, including engineering targeted herbicide or heavy metal tolerance and improving phytoremediation strategies.

In summary, ABCC1 orthologs from several agronomically relevant plant species were identified, and their biochemical properties were compared. Sequence and structural analyses indicate that these transporters share conserved features, including highly similar TMDs and NBDs, a bipartite substrate-binding pocket, and a degenerate NBS, and exhibit comparable in vitro ATPase activity. ZmABCC1 was further characterized and showed overlapping substrate specificity with AtABCC1. Collectively, these findings extend functional characterization of ABCC1 beyond the model species Arabidopsis and provide a framework for future studies aimed at improving crop performance and phytoremediation strategies.

## Methods

### Phylogenetic analysis of plant ABCC1

To identify ABC transporter genes across selected plant species, genomic analyses were performed using the Phytozome database (Goodstein et al., 2012). Primary amino acid sequences of all *Arabidopsis thaliana* ABC transporters were first retrieved from UniProt (The UniProt Consortium et al., 2023). These sequences (130 Arabidopsis ABC transporter genes) were used as queries in “search by BLAST” against each target plant genome within Phytozome using an E-value threshold of 1 x 10^−30^. Resulting hits were manually sorted to remove duplicate entries and alternative isoforms, generating a non-redundant list of ABC transporters for each species. This procedure was applied to *Zea mays* (maize), *Oryza sativa* (rice), *Sorghum bicolor* (sorghum), *Vigna unguiculata* (cowpea), *Phaseolus acutifolius* (tepary bean), *Glycine max* (soybean), and *Paspalum vaginatum* (seashore paspalum).

All identified plant ABC transporter sequences were subjected to phylogenetic analysis using Clustal Omega for multiple sequence alignment. The dataset also included all human ABC transporters, two additional plant ABCC1 homologs from *Phoenix dactylifera* (date palm) and *Carnegiea gigantea* (saguaro cactus), and orthologous ABCC1 proteins from *Saccharomyces cerevisiae* (Ycf1) and *Bos taurus* (BtABCC1). ABCC1 homologs were further analyzed using NCBI BLAST to calculate pairwise sequence identities relative to AtABCC1. Conserved residues within the predicted substrate-binding region of plant ABCC1 were identified from sequence alignments generated in MEGA12 and compared to the resolved substrate-binding site of bovine MRP1 (Johnson and Chen, 2017).

### Expression screening using Fluorescence Size Exclusion Chromatography

All genes were synthesized and cloned into the pFastBacMam expression vector with C-terminal Green Fluorescent Protein (GFP). Expression screening was performed in HEK293F cells grown in Freestyle 293 Expression Medium (Gibco) supplemented with 5% fetal bovine serum at 37 °C in 6-well plates. Transient transfection of adherent cells was carried out using 1 µg of ABCC1 plasmid DNA and 6 µg of linear polyethyleneimine per well. After 24 hours, valproic acid was added to a final concentration of 4.4 mM, and cells were harvested 72 hours post-transfection. Cell pellets were solubilized in buffer containing 50 mM Tris-HCl (pH 7.4), 150 mM NaCl, 1% (w/v) n-dodecyl-β-D-maltoside (DDM), and 0.2% (w/v) cholesteryl hemisuccinate (CHS) for 1 hour at 4 °C. Insoluble material was removed by centrifugation at 17,000 x g for 30 minutes, and the resulting crude lysate was analyzed by fluorescence-detection size-exclusion chromatography (FSEC) using an Agilent 1260 Infinity II HPLC system equipped with a fluorescence detector.

### Expression and Purification of Plant ABCC1

For large-scale expression and purification, the ABCC1 gene was cloned into pFastBac1 with C-terminal maltose binding protein (MBP) tag. This plasmid was transformed into DH10Bac cells to generate recombinant bacmid plasmid, and purified plasmid was introduced into Sf9 insect cells using transient transfection for baculovirus production. The generated baculovirus was amplified to P3 in Sf9 cells. For protein expression, the generated P3 virus was introduced (2% (v/v)) to Sf9 cells maintained in ESF 921 (Expression Systems) insect cell culture media with 2% (v/v) fetal bovine serum (FBS) at ∼2 x 10^6^ cells/mL cell density. After 72-hours of protein expression at 27 °C, cells were harvested by centrifuging at 1000 x g for 15 minutes and the cell pellet was flash frozen and stored at -80 °C until use.

Plant ABCC1 was purified by resuspending the cell pellet in a buffer containing 50 mM Tris-HCl (pH 7.4), 150 mM NaCl, 1% DDM, 0.2% CHS, and protease inhibitors including 2 µg/ mL Aprotinin, 10 µM Leupeptin, and 1 µM Pepstatin, and incubated for 1 hour at 4 °C. Then, the sample was centrifuged for 1 hour at 17000 x g at 4 °C to remove the insoluble material and the resulting supernatant was applied to a column with amylose affinity resin and incubated for 1 hour at 4 °C. The supernatant was then removed by gravity flow, and the resin was washed with a buffer containing 50 mM Tris-HCl (pH 7.4), 150 mM NaCl, 0.05% DDM, and 0.01% CHS. The protein was eluted with a buffer containing 50 mM Tris-HCl (pH 7.4), 150 mM NaCl, 0.05% DDM, 0.01% CHS and 10 mM maltose monohydrate. If MBP tag was to be removed, purified protein was incubated with MBP tagged TEV protease (TEV-MBP) in a ratio of 1: 5 (TEV protease: ABCC1-MBP) overnight at 4 °C. The next day, after performing six subsequent concentration-dilution steps to reduce the maltose monohydrate amount in the sample using a 100 kDa concentrator, the sample was applied to the second amylose affinity column to remove MBP tag and TEV-MBP protease. After 1 hour incubation, ABCC1 protein was collected in the flow-through of the column run and the sample was concentrated to < 500 µL. The digested sample was further purified with size exclusion chromatography (SEC) using Superose 6 gel filtration column with a buffer containing 50 mM Tris-HCl (pH 7.4), 150 mM NaCl, 0.05% DDM, and 0.01% CHS. Elution fractions containing purified ABCC1 were pooled and used for functional assays. Further, to verify the presence of purified plant ABCC1 protein, SDS-PAGE gel electrophoresis was conducted.

### ATPase activity assays

ATP hydrolysis (ATPase) activity assays were conducted in 96-well plates. ATPase activity was measured by detecting the release of inorganic phosphate through colorimetric assay. The amount of hydrolyzed phosphate was determined by comparison to a standard curve of potassium phosphate (KH_2_PO_4_). After performing initial assay condition optimizations (incubation time, incubation temperature and protein amount per reaction), all the assays were in a total reaction volume of 50 µL with 2 µg of purified ABCC1 protein, 5 mM ATP, 6 mM Mg^2+^, 50 mM Tris-HCl (pH 7.4), 150 mM NaCl, 0.05% DDM, and 0.01% CHS at 37 °C with 1 hour incubation time. Substrate concentration for all assays was 400 µM unless otherwise indicated. Vanadate inhibition was investigated with the addition of 2 mM sodium vanadate. All the assays were initialized by adding the protein, and after 1 hour reaction time, 50 µL of 12% (w/v) sodium dodecyl sulfate (SDS) was used to terminate the reaction and 100 µL from a mixture of 12% (w/v) ascorbic acid and 2% (w/v) ammonium molybdate (1:1) was used for color development. Absorbance was measured at 850 nm wavelength using a microplate reader. Each assay was conducted in triplicate, and the obtained data was analyzed using Microsoft Excel and GraphPad Prism software and presented as the average values ± standard deviation.

## Acknowledgements

Research reported in this publication was supported by the National Institute of General Medical Sciences of the National Institutes of Health under Award Number R35GM166048. The content is solely the responsibility of the authors and does not necessarily represent the official views of the National Institutes of Health.

## Data availability

Data will be made available on request.

