## Supplementary Figures for "Plant ABCC1 orthologs from agronomically relevant species reveal conserved principles of detoxification transport"

**Table of Contents**

Supplementary Figure 1. Phylogenetic analysis of plant ABC transporters…………………… 3

Supplementary Figure 2. Expression Screening and purification of plant ABCC1 homologs… 4

Supplementary Figure 3. ATPase Activity of ZmABCC1……………………………………… 5

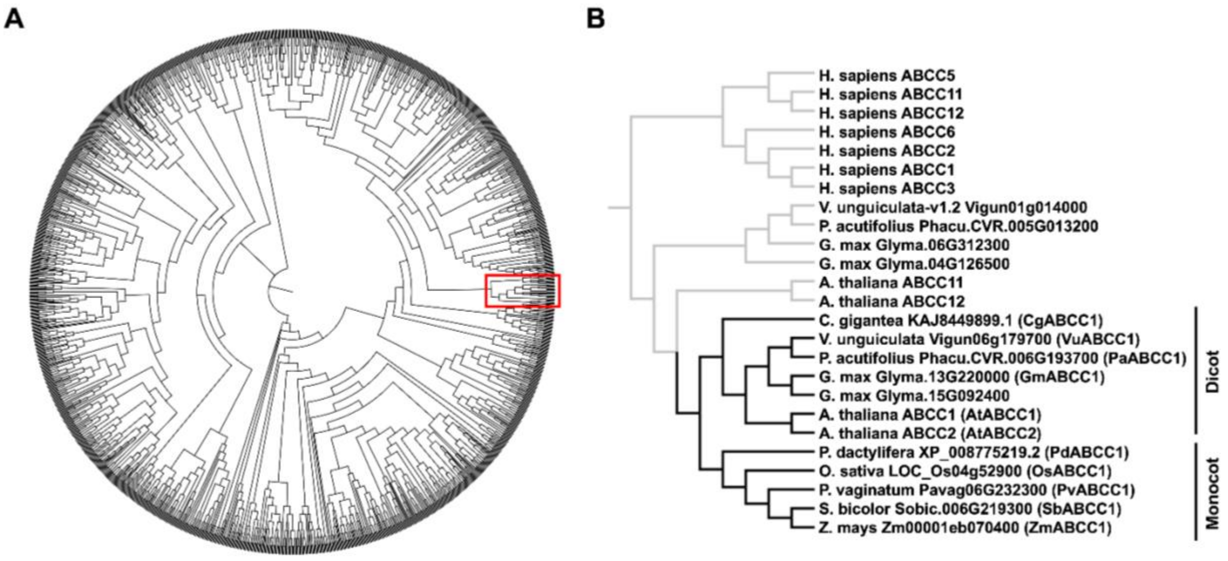


**Supplementary Figure 1. Phylogenetic analysis of plant ABC transporters. A.** Phylogenetic tree constructed from complete ABC transporter dataset. All of the full-length ABC transporter sequences from each plant species were retrieved from the Phytozome database and used to construct a phylogenetic tree. Plant species included *Arabidopsis thaliana*, *Zea mays* (maize), *Oryza sativa* (rice), *Sorghum bicolor* (sorghum), *Vigna unguiculata* (cowpea), *Phaseolus acutifolius* (tepary bean), *Glycine max* (soybean), and *Paspalum vaginatum* (seashore paspalum). In addition, all the ABC transporters from *Homo sapiens* (human) were included, along with ABCC1 homologs from *Bos taurus* (bovine), *Saccharomyces cerevisiae* (yeast), *Phoenix dactylifera* (date palm) and *Carnegiea gigantea* (saguaro cactus). The red box indicates position of ABCC1 homologs, which are expanded in panel (B). Due to the size of the phylogenetic tree, protein names are not displayed. **B.** Expanded view of the ABCC1 clade. Direct ABCC1 orthologs are shown in black bars, whereas related paralogs within the same branch are indicated with grey bars. Only orthologs indicated by black bars were selected for further analysis.

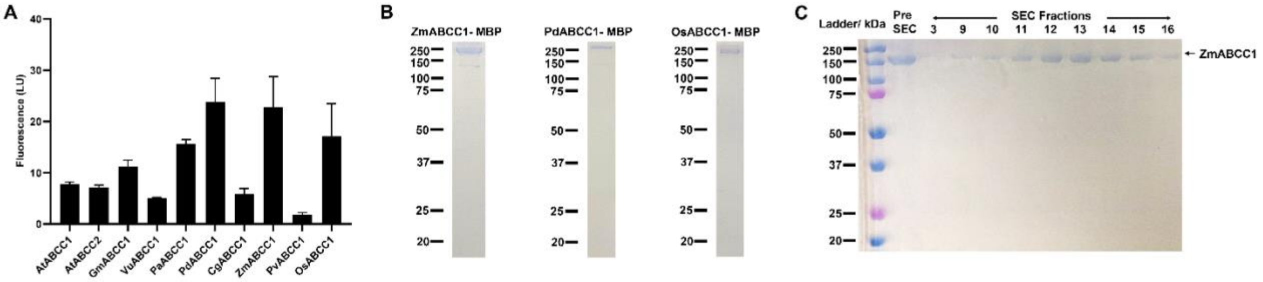


**Supplementary Figure 2. Expression Screening and purification of plant ABCC1 homologs.** **A.** Quantitative analysis of expression level of plant ABCC1 homologs using FSEC. Protein expression screening was performed in duplicates and the maximum values of FSEC chromatograms of the ABC transporter peak were averaged and graphed. Representative chromatograms from one replicate are shown in Figure 4A. **B.** SDS-PAGE gel of purified C-terminal MBP tagged plant ABCC1 orthologs, including ZmABCC1, PdABCC1, and OsABCC1. The samples were used for the ATPase assays shown in Figure 4B-C. **C.** SDS-PAGE gel of the fractions collected from size exclusion chromatography of MBP cleaved, purified ZmABCC1. The corresponding chromatogram is shown in Figure 4D.

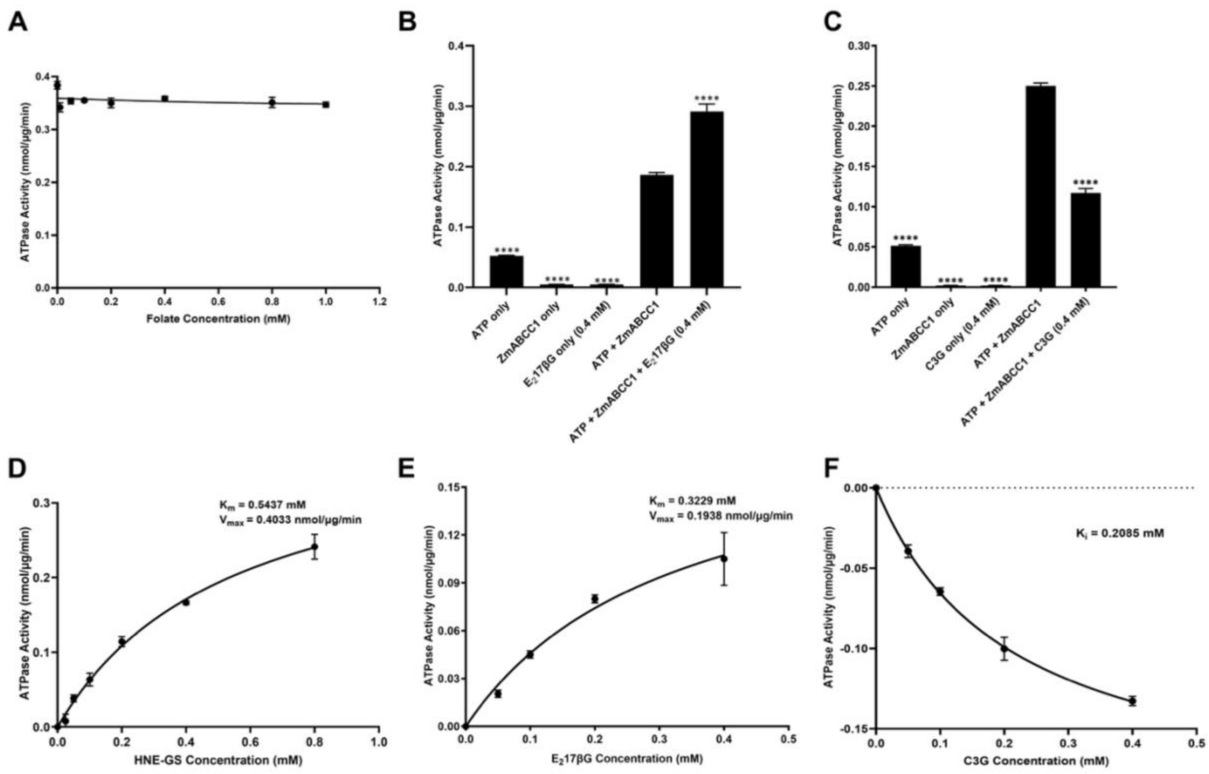


**Supplementary Figure 3. ATPase Activity of ZmABCC1. A.** ATPase activity of purified ZmABCC1 with Folate up to 1 mM concentration. No change in ATPase was observed for any of the concentrations tested. **B-C.** ATPase activity of purified ZmABCC1 with the control experiments in the presence of $E_2$17βG (**B**) and C3G (**C**). **D-F**. Saturation curves of HNE-GS (**D**), $E_2$17βG (**E**), and C3G (**F**). Data in D-F is the identical data to that shown in Figure 6, but all data points have basal ATPase subtracted to calculate kinetic parameters. Statistical significance of the data in panel A was determined by one-way ANOVA followed by Dunnett's multiple comparison test (ns, $P \geq 0.05$ (not statistically significant); *, $P < 0.05$; **, $P < 0.01$; ***, $P < 0.001$; ****, $P < 0.0001$).
